# A Method for Electrical Stimulus Artifact Removal Exploiting Neural Refractoriness: Validation by Contrasting Cathodic and Anodic Stimulation

**DOI:** 10.1101/2024.10.06.616879

**Authors:** A. Nakhmani, J. Block, M. Awad, J. Olson, R. Smith, J.N. Bentley, M. Holland, S.A. Brinkerhoff, C. Gonzalez, M. Moffitt, H. Walker

## Abstract

**Objective:** To present a novel method for removing stimulus transient that exploits the absolute refractory period of electrically excitable neural tissues.

**Background:** Electrical stimulation often generates significant signal artifacts that can obscure important physiological signals. Removal of the artifact and understanding latent information from these signals could provide objective measures of circuit engagement, potentially driving advancements in neuromodulation research and therapies.

**Methods:** We conducted intracranial physiology studies on five consecutive patients with Parkinson’s disease who underwent deep brain stimulation (DBS) surgery as part of their routine care. Monopolar stimuli (either cathodic or anodic) were delivered in pairs through the DBS electrode across a range of inter-stimulus intervals. Recordings from adjacent unused electrode contacts used broadband sampling and precise synchronization to generate a robust template for the stimulus transient during the absolute refractory period. These templates of stimulus transient were then subtracted from recordings at different intervals to extract and analyze the residual neural potentials.

**Results:** After artifact removal, the residual signals exhibited absolute and relative refractory periods with timing indicative of neural activity. Cathodic and anodic DBS pulses generated distinct patterns of local tissue activation, showing phase independence from the prior stimulus. The earliest detectable neural responses occurred at short peak latencies (ranging from 0.19 to 0.38 ms post-stimulus) and were completely or partially obscured by the stimulus artifact prior to removal. Cathodic stimuli produced stronger local tissue responses than anodic stimuli, aligning with clinical observations of lower activation thresholds for cathodic stimulation. However, cathodic and anodic pulses induced artifact patterns that were equivalent but opposite.

**Interpretation:** The proposed artifact removal technique enhances prior approaches by allowing direct measurement of local tissue responses without requirements for stimulus polarity reversal, template scaling, or specialized filters. This approach could be integrated into future neuromodulation systems to visualize stimulus-evoked neural potentials that would otherwise be obscured by stimulus artifacts.

## Introduction

Deep brain stimulation (DBS) is a widely used therapy that alters neural activity in patients with neurological and psychiatric diseases (1). Beyond DBS, both implanted and external electrical stimulation devices are used across various medical fields, including rehabilitation (2), pain management (3), cardiology (4), gastroenterology (5), urology (6), and others. In research, these devices play crucial roles in probing circuit function to identify physiological mechanisms. Whether for therapy or research, comprehensive visualization of stimulus-evoked activity is essential, as it offers insights into circuit engagement, which could potentially serve as a proxy for dose. For clinical applications, these measurements could guide targeting and programming, and inform more sophisticated lead designs or adaptive feedback systems (7–10).

In the context of DBS for movement disorders, electrophysiological biomarkers have emerged as candidate tools to optimize therapy. These biomarkers fall into two broad categories – spontaneous / task-related field potentials (11) and stimulus-evoked activity (12–15). While beta oscillations (∼13-30 Hz) have shown promise, (16,17) they are not always present and are vulnerable to interference from cardiopulmonary activity, neck movements, and other sources of noise. In contrast, stimulus-evoked potentials display larger amplitudes and are more resilient to noise because of signal averaging (18,19). Consequently, interest in DBS-evoked potentials within the basal ganglia, thalamus, and cerebral cortex is growing. Specifically, stimulation in the subthalamic nucleus (STN) and globus pallidus interna (GPi) in PD and dystonia patients evokes resonant neural activity (ERNA), a fast, tapering oscillatory potential within the basal ganglia (14,15). ERNA amplitude correlates with therapeutic efficacy (15,20–23) and also shows short-term neuroplasticity (12), suggesting a functional interaction with local basal ganglia circuits. ERNA might therefore inform the selection of stimulation parameters or provide dynamic feedback for real-time stimulation adjustments (14).

Defining stimulation dose is a persistent challenge in neuromodulation, particularly as it relates to brain circuit physiology. One problem is that electrical stimuli generate large, high-frequency transients that obscure fast brain dynamics. Various methods attempt to address this in the context of DBS. The most common approach is to ignore the fastest components of the signals, which is suitable for local field potential studies that focus on spectral content beneath clinical DBS frequencies (<100 Hz). Other methods blank or interpolate the artifact over a short duration (24,25) or else use template subtraction (26–30), independent components analysis (31), specialized filters (30,32), shape estimation (33), or non-causal Weiner filtering (34). While partially effective, these methods are difficult to validate and prone to various forms of information loss and/or signal distortion, limiting their use to research applications.

Our prior work used stimulus polarity reversal to remove the artifact by phase cancellation (12,13,35,36). Summing pairs of DBS-evoked potentials arising from opposite stimulus polarities yields cancels or minimizes the artifact and amplifies the brain response of interest, enabling direct visualization of novel short-latency cortical and subcortical potentials that were otherwise obscured (12,13,35). However, this method has its own limitations: phase cancellation is not always perfect, the assumption that polarity does not affect the brain response is often incorrect, and the composite waveform is derived from two sets of stimulation parameters, complicating dose estimation. Additionally, whether the earliest components of the evoked responses (R1) arise from brain activity is still a matter of debate (12,13,35).

Here we propose a novel approach to stimulus artifact removal that exploits the absolute refractory period of neural tissues, building on prior work from the peripheral nervous system (37). We validate this approach experimentally by demonstrating neural refractoriness and by contrasting basal ganglia field potential responses to cathodic and anodic stimuli. The method relies on generating a template waveform for the stimulus artifact during the absolute refractory period using paired stimulus pulses. Subtracting this template from other stimulation events reveals neural activity that would otherwise be obscured by the artifact. This procedure, or its extensions, could eventually be embedded into device hardware or firmware to enable real-time visualization of fast neural dynamics and provide new insights into dose or circuit engagement by neuromodulation therapies.

## Materials and Methods

### Participants

This project received prior approval from the UAB Institutional Review Board, and all participants provided written informed consent prior to enrollment. All participants were diagnosed with Parkinson’s disease (PD) by a movement disorders neurologist and underwent sub-thalamic nucleus DBS as part of routine care.

*DBS surgery, behavioral assessment, and clinical stimulation*. Surgeries were performed on awake patients in the ‘off’ medication state using frame-based stereotaxy. Prior to the research, we administered local anesthesia and intravenous midazolam at a dosage of 1-2 mg, approximately 1-3 hours before frame placement. Abbott 8-contact ‘1-3-3-1’ directional DBS leads were implanted with targeting guided by multi-pass microelectrode recordings, micro- and macro-stimulation, O-arm 2 CT imaging, and real-time anatomic reconstructions using BrainLab software. Motor symptoms were assessed intraoperatively at three time points: baseline (pre-DBS), immediately after lead implant with the device off, and during trial stimulation with DBS. Assessments utilized sub-scores from the Unified Parkinson’s Disease Rating Scale (UPDRS) part 3 for the contralateral upper and lower extremities. Trial DBS parameters used cathodic stimulation and were optimized to enhance motor improvement beyond microlesion effects, with a pulse width of 60 μs and a frequency of 130 or 160 Hz, per standard clinical practice.

### Experimental stimulation and signal acquisition

An external pulse generator (STG4002, Multi-Channel Systems, Reutlingen, Germany) delivered pairs of monophasic rectangular wave pulses through electrodes on the implanted DBS electrode array. Monopolar cathodic and anodic stimuli were delivered through one DBS electrode using cathodic DBS parameters that displayed clinical efficacy during trial stimulation. Pairs of identical DBS pulses were delivered across 16 unique inter-stimulus intervals ranging from 0.3 to 16 ms in randomized blocks. To ensure precise charge balance, we delivered an active charge recovery pulse around 20 ms (jittered) after each pair of stimuli. Charge density never exceeded the FDA-recommended limit of 30 µC/cm^2^/phase, and the experimental stimulation was physically imperceptible in all participants. Pauses between stimulus pairs were randomly uniformly distributed between 50 to 60 ms, yielding a mean pause of 55 ms. A transistor-transistor logic (TTL) signal was generated for each stimulus, providing sync precision of 10 μs. Non-essential electrical equipment was deactivated, and unsaturated local field potentials were recorded using an actiCHamp amplifier (Brain Vision LLC, Morrisville, NC). We established a ground on the scalp outside the sterile field and sampled at 100 kHz with an analog low pass filter of 7.6 kHz.

### Digital signal processing

We introduce a novel method to isolate the neural response to a single set of DBS parameters using pairs of monopolar stimuli. The recorded signals were analyzed with EEGLAB and custom routines in MATLAB (Mathworks, Natick, MA). To enhance the isolation of signals near the stimulation site, the recordings from the DBS electrode were re-referenced to a bipolar montage. During paired-pulse stimulation, our primary interest is the neural response to the second stimulus (test stimulus). To isolate this response, the activity from the first stimulus (conditioning stimulus) was removed using template subtraction (12). At very short interstimulus intervals (<0.5 ms), the test stimulus is divorced from local neural activity because the stimulus was initiated within the absolute refractory period. We leveraged this phenomenon to generate a second template waveform representing the components of the potential associated with the stimulus alone (see **Schema**). This allows the isolation of local brain responses, even those at short latencies that would otherwise be obscured by the stimulus artifact. Although the focus of this study was monopolar stimuli, similar methods can be applied to s and more complex paradigms and bipolar stimulation.

To describe our approach more specifically, assume that the response to a given stimulus pulse is *A*(*t*) + *B*(*t*) where *A*(*t*) is stimulus artifact, and *B*(*t*) is the brain response. When in the absolute refractory period, the second stimulus pulse (the test stimulus) is delayed by a short time *A*(*t* − τ), where τ is the time delay, with no associated brain response because of neural refractoriness. The total response to the pair of stimulus pulses is *A*(*t*) + *B*(*t*) + *A*(*t* − τ). Therefore, if we subtract the single pulse response from the paired pulse response, the remainder will consist of a template of the artifact at a time delay τ. We can compute the template *A*(*t*) at time *t* by shifting *A*(*t* − τ) backward in time by the delay τ. Finally, we subtract the template for *A*(*t*) from the conditioning and testing (or single pulse) responses to isolate the neural response alone (final row of **Schema**).

### Characterization of evoked potentials

Our primary goal was to characterize subcortical local field potentials within the first 10 ms after the stimulus onset, including short latency R1 potentials in close temporal proximity to the stimulus and later oscillatory activity (ERNA). All responses were aligned to the test stimulus using TTL sync markers, and the signal traces were color-coded based on paired pulse interval (**Figure 1**). To compare responses at different interstimulus intervals within each participant, we normalized response magnitude to the template waveform of the conditioning stimulus, allowing calculation of the paired pulse ratio. Paired pulse ratio is defined as the test stimulus response magnitude divided by the conditioning stimulus response magnitude. A ratio of 1 indicates that the test stimulus recruited a response of identical magnitude to the conditioning stimulus, whereas a ratio of 0 indicates no neural recruitment by the second stimulus. We defined the onset of the absolute refractory period as the first interstimulus interval among 3 consecutive paired pulse ratios of <0.2 to account for the noise floor. The relative refractory period was chosen as the first interstimulus interval among 3 consecutive paired pulse ratios of <0.98.

**FIGURE 1.**
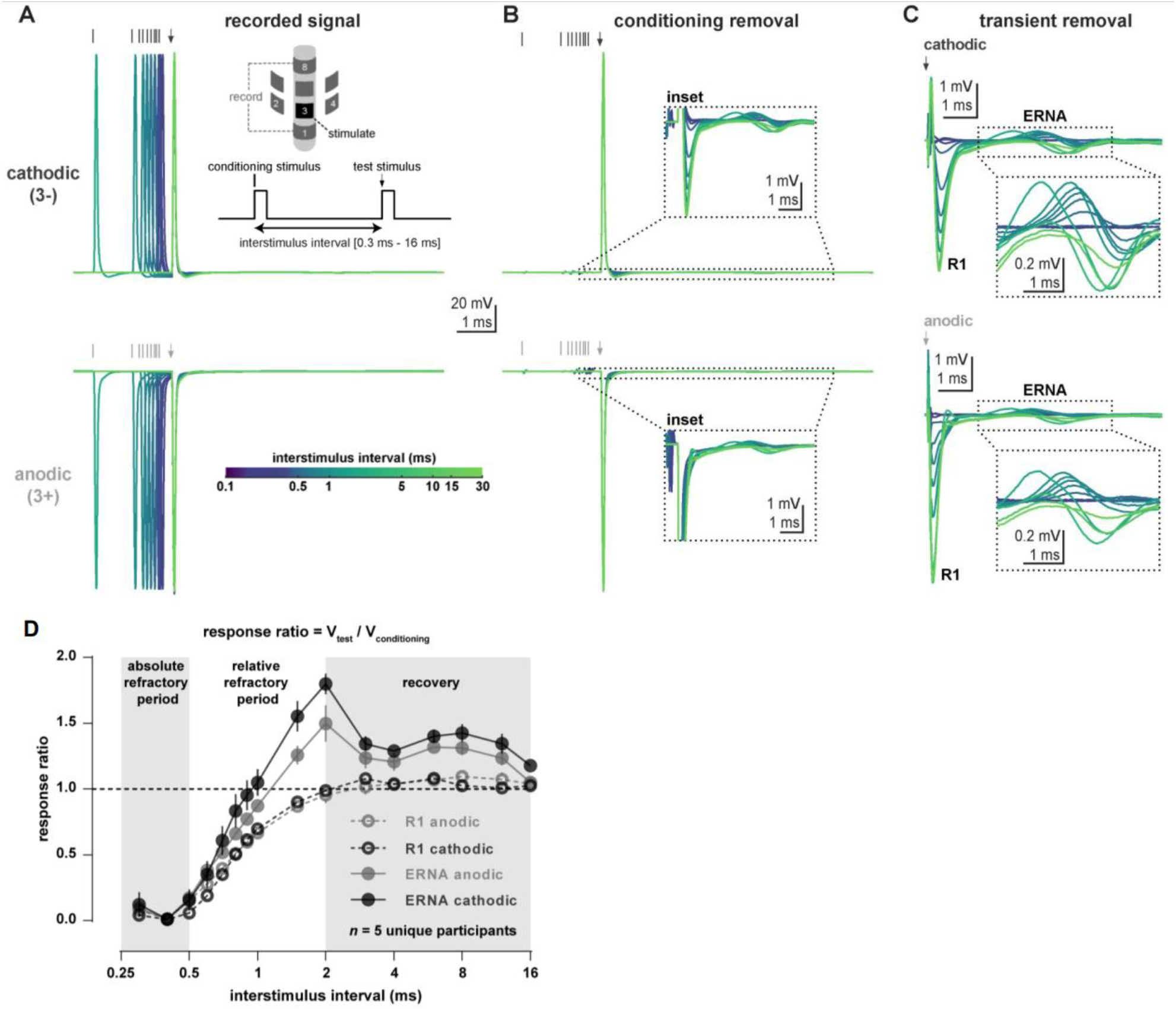
Monopolar cathodic and anodic DBS pulses elicit local tissue responses across multiple time scales that display refractoriness and short-term plasticity. Panels (A)-(C) show results for one representative participant, and panel (D) shows aggregate average results for all participants. **(A)** *Pairs of DBS pulses were delivered from a directional electrode across a range of interstimulus intervals, and field potential responses were sampled from a bipolar pairing of the outer ring contacts*. **(B)** *Stimulus artifact and neural responses associated with the first pulse (conditioning stimulus) responses were first removed by template subtraction*. ***Insets*** *show that stimulation elicits neural activation that is fully or partially embedded in the stimulus artifact from the second pulse (test stimulus), regardless of cathodic or anodic stimulus polarity*. **(C)** *At very short interstimulus intervals (∼0*.*3 ms), the electrical stimulus transient is divorced from neural activation because of the absolute refractory period. We subtract a template of the stimulus artifact alone from the test stimuli across the other paired pulse intervals. This reveals potentials that were previously obscured by artifact*. **(D)** *Paired cathodic and anodic pulses both elicit responses that display absolute and relative refractory periods. The later evoked resonant neural response (ERNA) displays paired pulse facilitation at specific interstimulus intervals in response to both anodic and cathodic stimulation*.

**SCHEMA.**
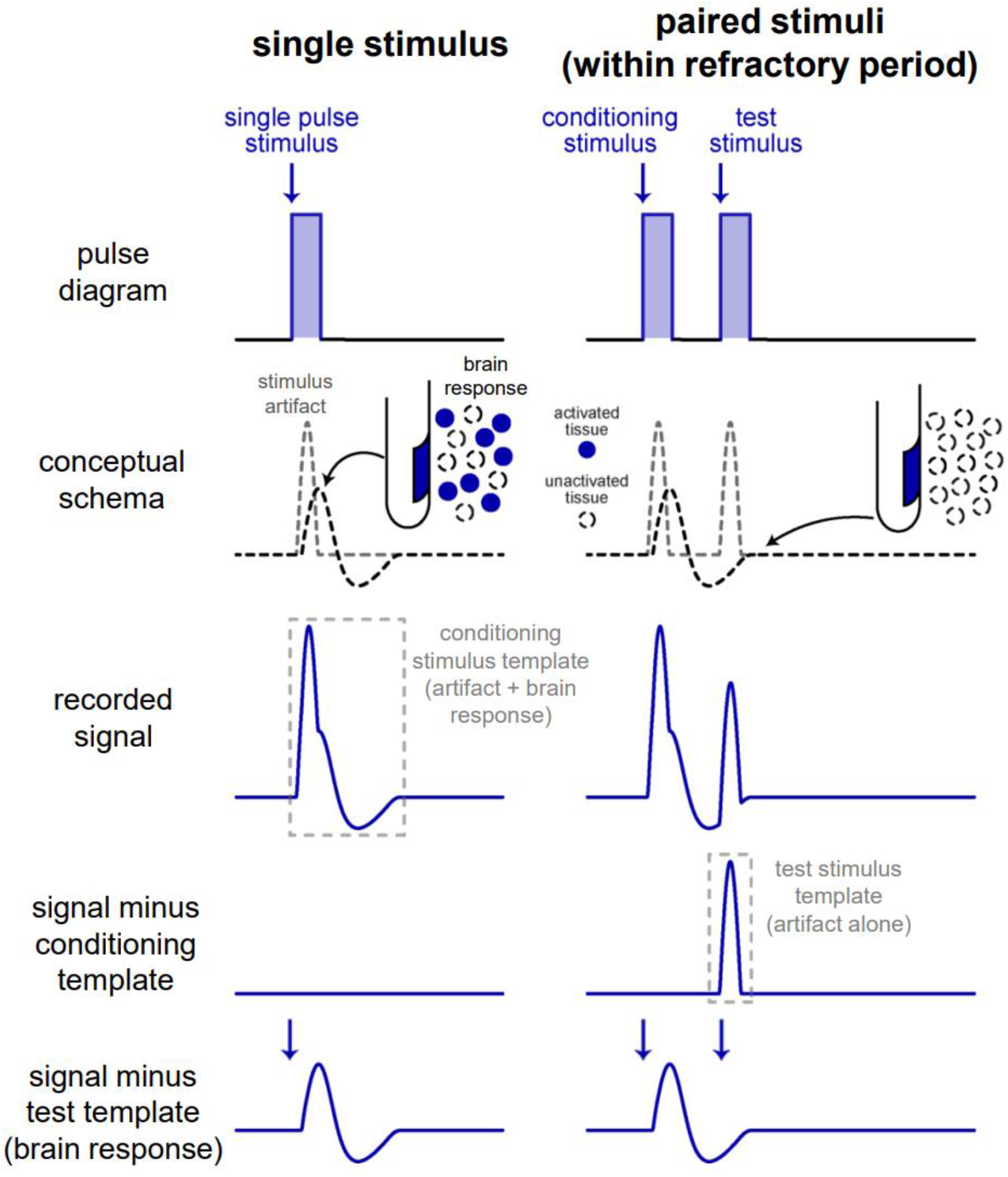
Stimulus artifact removal method exploiting the absolute refractory period of electrically excitable tissues. Electrical stimulus pulses generate large signal transients (‘artifacts’) that obscure the neural response to stimulation. Equivalent (or lesser) stimuli within the absolute refractory period cannot recruit neural elements, yielding a template waveform for the stimulus artifact alone. Subtraction of this template allows isolation of the neural response to any DBS pulse that uses identical stimulation parameters.

## Results

*Demographics and clinical data*. We studied intracranial signals from basal ganglia in five consecutive patients with Parkinson’s disease who underwent DBS surgery as part of routine care. The average (SD) participant age was 63.6 (14.7) years, and the duration of disease on the day of surgery was 7.1 (1.5) years. All were white males who underwent left-hemisphere STN implants.

*DBS-evoked subcortical electrophysiology: Response dynamics and plasticity*. Monopolar stimuli were delivered from a directional contact within the STN, and recordings were obtained from the outer ring contacts. Despite the physical proximity of the monopolar stimulus and recording sites, artifact template subtraction revealed both R1 and the later ERNA responses in 5/5 participants. Cathodic and anodic stimuli both elicited R1 at peak latencies of 0.3 ± 0.1 ms and amplitudes of 1.1 ± 0.9 and 1.0 ± 0.8 mV, respectively.

Resonant oscillatory activity (denoted ERNA) in the STN circuit was recorded as well in all 5 patients. **Figure 1** shows that our method could be used to remove the stimulus artifact regardless of the stimulus polarity. The subject shown in this figure represents the ideal case where the earliest response (R1) to electrical stimulation is not completely obstructed by the stimulus transient. Using our method did not alter this R1 response, which suggests that our method extracts the true transient signal produced by the stimulus pulse. The obtained paired-pulse ratio plot in Panel D is very similar to previously published manuscripts in this field (12). This also adds more support to our method and provides further evidence of the successful stimulus artifact removal. Panels A-C of **Figure 1** show how our method was applied separately to cathodic and anodic stimulation responses using independent templates. **Figure 1** also shows that this method works regardless of the stimulus polarity. The artifact was removed for both cathodic and anodic stimulation which helps in the comparison of local tissue activation between the two stimulation polarities. Note that the stimulus artifacts for cathodic and anodic stimulation (**Figure 1 A,B**) are of opposite direction, as expected, but that the neural responses are in the same direction, providing further support for the method. The artifact removal method was applied to 4 more subjects, and it showed similar results in that it removed the stimulus artifact from the neural response.

In **Figure 2**, we show the relation between the area of R1 (left panel) for anodic and cathodic stimulation and similarly the relation between the area under the ERNA oscillation (right panel) for anodic and cathodic stimulation. This information helps compare the necessary dose for the different polarities of stimulation. First, the area for the R1 response was scaled between 0 and 1 to reduce the effects of impedance differences across patients. Similarly, the area under the rectified ERNA was normalized between 0 and 1. The dashed unity line corresponds to an equal dose response.

**FIGURE 2.**
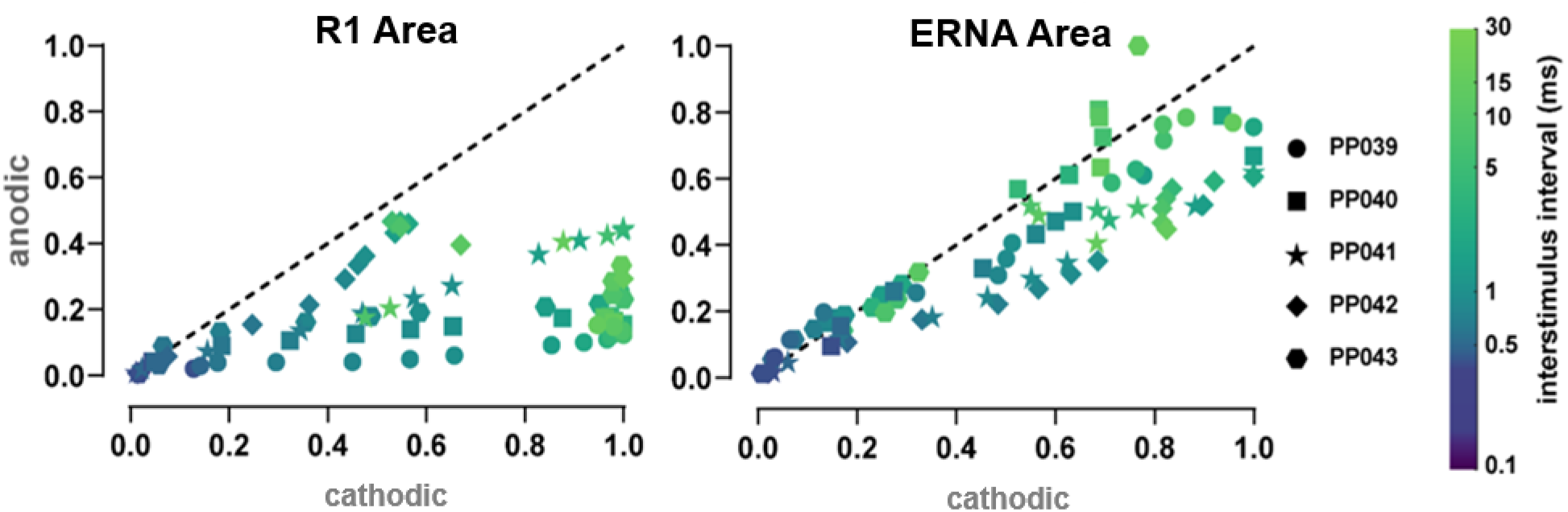
Monopolar cathodic and anodic pulses elicit distinct local neural responses. **(A)** Scatter plots and diagonal unity lines, with response area normalized to the largest potential within each participant. Cathodic stimuli elicit early (R1) and later responses (ERNA) with larger integrated area than anodic stimuli within the same participant.

As can be seen in **Figure 2**, within the absolute refractory period both anodic and cathodic response areas are very similar and have negligible magnitudes, as expected. For longer interstimulus intervals of paired pulses, the R1 areas (left panel) always have smaller magnitudes for anodic stimulation versus cathodic and they cap at around half the size of the cathodic pulse response area. For ERNA responses (right panel) to paired pulses with interstimulus intervals longer than the absolute refractory period, the responses are also typically smaller for anodic stimulation, but not to the same degree as R1.

## Discussion

Our findings highlight an innovative approach to visualize short latency stimulus-evoked field potentials that were previously obscured by the electrical stimulus artifact from DBS. The proposed method exploits refractoriness, a fundamental physiological property of electrically excitable tissues, to isolate and remove components of the potential that arise from stimulus artifacts and not neural activity. Furthermore, contrasts between responses to monopolar cathodic and anodic stimuli provide additional validation for the neural origin of the earliest components of these potentials.

Stimulus artifacts challenge research and translational applications in neuromodulation because they obscure the fast dynamics of the underlying neural activity, especially when recording and stimulation sites are close to one another. This issue is particularly prevalent in DBS research where therapeutic stimulation consists of brief pulses delivered at relatively high frequencies (>100 Hz). Consequently, most studies focus on lower frequency components of the signal or more delayed responses between consecutive pulses. Our methods or their extensions could provide a more objective basis for measuring stimulation dose or circuit engagement within an individual, with the potential to advance both research and clinical applications. Furthermore, they could be applied in other clinical domains where electrical stimulation interacts with excitable tissues in the central, peripheral, autonomic, or cardiac systems.

Multiple observations substantiate our claim that the residual signals after artifact removal arise from neural activity. First, all potentials, including the shortest latency components that would otherwise be obscured by stimulus artifact, exhibit absolute and relative refractory periods at paired pulse intervals compatible with neural activity (38,39). Second, anodic and cathodic stimuli yield artifacts with opposite polarities, yet the subsequent potentials after artifact removal do not display phase reversals. This indicates that the stimulus artifact and the later potentials (R1, ERNA) arise from distinct sources. Finally, cathodic stimuli elicit larger potentials than otherwise identical anodic stimuli, consistent with observations spanning decades that cathodic electrical fields activate neural elements at lower amplitude thresholds. Taken together, these observations provide independent and compelling evidence that both R1 and ERNA arise from stimulus-evoked neural activity.

Our findings have several implications for the DBS mechanism of action. First, R1 occurs at 0.3±0.1 ms peak latency and is typically obscured by stimulus artifact. This potential likely represents the earliest detectable interaction between a given DBS pulse and the local tissue environment when recording from intracranial macroelectrodes. Second, R1 in the basal ganglia exhibited shorter peak latencies than similar recordings from the cortical surface (12,40) and the scalp (13,35,40). These longer delays at more distant recording sites are therefore most consistent with retrograde (antidromic) propagation via the cortico-subthalamic (hyper-direct) pathway rather than volume conduction of the subcortical potentials. Third, our data show that interstimulus intervals corresponding to therapeutic DBS frequencies (∼6-10 ms) are well outside the absolute and relative refractory periods of the neural elements. This suggests that the immediate effect of single and paired DBS pulses in humans involves depolarization and synchronization of surrounding neural elements to the stimulation frequency. Future studies should investigate the temporal evolution of these responses during bursts and longer stimulus trains.

Our research also sheds light on anodic stimulation, a relatively new option for DBS therapy. Keeping all other stimulation parameters constant, we found greater local tissue activation with cathodic versus anodic DBS pulses. Neuromodulation pioneer James Ranck proposed that stimulus polarity differentially alters the recruitment of specific neural elements. He hypothesized that cathodic stimuli display lower excitation thresholds for axons and fibers of passage, whereas anodic stimuli interact primarily with local cell bodies (41), particularly those with axons oriented away from the electrode. Computational modeling studies (42–46) have since supported these claims, although human work remains limited (47,48). An early clinical study confirms different excitation thresholds for DBS side effects by polarity, consistent with Ranck’s idea that anodic and cathodic stimuli activate distinct but overlapping neuronal populations (47). This double-blind crossover pilot study also found more favorable clinical outcomes with anodic over conventional cathodic stimulation in PD patients without a predominant tremor. Our results in humans provide electrophysiological and mechanistic evidence to support these claims.

Several technical questions arise from our approach, as it employed a *post hoc* software-based implementation. The minimum sampling frequency and synchronization accuracy to yield acceptable results is unclear. In theory, faster sampling and/or more precise synchronization should improve artifact suppression because our stimuli were square wave pulses with infinite frequency content. Higher sampling frequencies, however, come at the costs of greater heat generation, battery usage, and memory requirements. While less a concern for an externalized decision-assistance tool or a discrete assay, this could present practical barriers to developing fully implanted, self-contained hardware for closed-loop stimulation. ERNA and other features in the signal, however, likely would not require such rapid sampling, an epoch-based approach to signal acquisition could significantly decrease sampling demand, as well. Hardware-based implementations could also be viable, as demonstrated by recordings from the median nerve in the periphery by McGill et al. (37). Finally, stimulus artifacts are typically large, therefore amplifier gain must be large enough to accommodate the full range of the entire stimulus transient and the anticipated brain response.

The proposed approach has several notable advantages. Importantly, essentially all prior studies do not recognize or evaluate physiological brain activity at these short latencies, because the responses are obscured by stimulus artifacts. These responses display large amplitudes and dose dependence and can occur regardless of whether local oscillatory activity follows (12). Another feature is that this method isolates the local neural response to a single set of stimulation parameters. Prior work using phase cancellation with stimuli of opposite polarities yields an aggregate response waveform that overestimates tissue activation from either polarity alone. Another assumption of the phase cancellation method is that stimuli of opposite polarities yield similar neural responses, but here we show that anodic and cathodic DBS pulses elicit different event-related local potentials in the basal ganglia. Another interesting facet of this work is that artifact reduction performed well using monopolar stimuli. Monopolar stimuli typically yield much larger stimulus artifacts than bipolar stimuli, so this signal processing algorithm conceivably might perform better with bipolar stimuli. Finally, while we have focused on DBS as a test case, same methods could be applied to better understand how other electrical stimulation modalities impact neural responses at both local and more distant recording sites.

These methods have potential limitations, as well. A possible barrier to clinical adoption is that broadband sampling increases power consumption, heat generation, and data storage requirements. As an initial implementation, these factors are less problematic with peripheral hardware, for instance as a decision assistance tool during targeting in the operating room. Fully implanted hardware with stimulation and sensing capabilities could first be implemented at discrete time points to inform open-loop stimulation (similar to an impedance test or a query of spontaneous field potentials in current clinical workflows). Adaptive stimulation using real-time recordings could benefit from algorithm optimization, rechargeable pulse generators, and other hardware advances. Additionally, continuous sampling is likely unnecessary for generating event-related potentials, once the time range of the signal of interest has been defined. Another important limitation is that we so far cannot accurately remove stimulation artifacts on a pulse-by-pulse basis, as evoked potential calculation typically requires dozens of stimulation events at a minimum. Clinical DBS however is typically delivered at relatively high frequencies (>100 Hz). As such, moving averages could yield feedback in fractions of a second, which is likely prompt enough for behavioral relevance in movement disorders applications. Finally, our study examined responses to single and paired DBS pulses at the STN target. Future work should examine how these responses evolve in response to bursts and continuous stimulation, and how response timing and morphology differ across different stimulation and recording sites.

## Conclusions

The proposed artifact removal method holds significant potential for various applications in both research and clinical domains related to DBS and other forms of neuromodulation. Our method provides a detailed visualization of how local tissues respond to stimulation, offering new insights into traditional ideas about how stimulation parameters influence dose or activate neural circuits. These findings could inform the parameterization of implanted or externalized commercial hardware with greater potential to personalize DBS therapy or to serve as feedback signals for closed-loop stimulation. More broadly, better artifact removal methods have the potential to enhance patient outcomes not only in the context of deep brain stimulation but also in other domains where electrical stimuli interact with excitable tissues.

## Acknowledgments

Research reported in this publication was supported by NINDS of the National Institutes of Health under award number UG3NS130202. RS was supported by the American Epilepsy Society Junior Investigator Award 1042632, CURE Epilepsy Foundation Taking Flight Award 1061181, and the ORAU Ralph E. Powe Faculty Enhancement Award. JB was supported by Boston Scientific. The content is solely the responsibility of the authors and does not necessarily represent the official views of the National Institutes of Health.

